# Stage-specific transcriptomes and DNA methylomes indicate an early and transient loss of transposon control in Arabidopsis shoot stem cells

**DOI:** 10.1101/430447

**Authors:** Ruben Gutzat, Klaus Rembart, Thomas Nussbaumer, Rahul Pisupati, Falko Hofmann, Gabriele Bradamante, Nina Daubel, Angelika Gaidora, Nicole Lettner, Mattia Donà, Magnus Nordborg, Michael Nodine, Ortrun Mittelsten Scheid

## Abstract

In contrast to animals, postembryonic development in plants is modular, and aerial organs originate from stem cells in the center of the shoot apical meristem (SAM) throughout life. Descendants of SAM stem cells in the subepidermal layer (L2) give also rise to male and female gametes (reviewed in ^1^) and are therefore considered primordial germ cells. In these cells, transmission of somatic mutations including virus and TE insertions must be avoided. Despite their essential role for plant development and intergenerational continuity, no comprehensive molecular analysis of SAM stem cells exists, due to their low number, deep embedding among non-stem cells, and difficult isolation. Here we present a comprehensive analysis of stage-specific gene expression and DNA methylation dynamics in Arabidopsis SAM stem cells. Stem cell expression signatures are mostly defined by development, but we also identified a core set of differentially expressed stemness genes. Surprisingly, vegetative SAM stem cells showed increased expression of transposable elements (TEs) relative to surrounding cells, despite high expression of genes connected to epigenetic silencing. We also find increasing methylation at CHG and a drop in CHH methylation at TEs before stem cells enter the reproductive lineage, indicating an onset of epigenetic reprogramming at an early stage. Transiently elevated TE expression is reminiscent of that in animal primordial germ cells (PGCs) ^2^ and demonstrates commonality of transposon biology. Our results connect SAM stem cells with germline development and transposon evolution and will allow future experiments to determine the degree of epigenetic heritability between generations.

---

In Arabidopsis, the SAM stem cell niche is marked by expression of *CLAVATA3 (CLV3)*. While other transcription factors, signaling molecules (including *CLV3*), and receptors (reviewed in ^3, 4^) are necessary for stem cell maintenance, our knowledge of the characteristics of “stemness” and the molecular signatures of plant stem cells remains limited. Previous studies have used mutants to increase CLV3-expressing cells, with associated phenotypic abnormalities, or used entire meristems including non-stem cells ^5–8^. To obtain information about gene expression and DNA methylation of pure SAM stem cell fractions, we generated Arabidopsis plants expressing a transcriptional fusion of the *CLV3* promoter ^9^ and mCherry-abelled histone H2B. Microscopic analysis demonstrated correct and specific expression of the p*CLV3:mCherry-H2B* marker in nuclei of ≈20-40 stem cells in 14-day-old seedlings (Fig. 1a). We applied fluorescence-activated nuclear sorting (FANS) ^10^ to nuclei isolated from tissue manually enriched for shoot apical meristems and collected mCherry-positive and-negative nuclei, with non-transgenic plants as controls (Fig. 1b, Fig. S1a, Table S1). Microscopic analysis confirmed that all sorted nuclei from the positive channel appeared intact and displayed red fluorescence, validating the purity of the fraction (Fig. S1b). The transcript level of endogenous *CLV3* was more than 1000-fold higher in mCherry-positive versus controls (Fig. 1c). RNA expression in sorted nuclei was highly correlated with RNA from tissue samples (Pearson correlation coefficient = 0.94; Fig. S1c), indicating that nuclear RNA represents the transcriptome of whole cells. Therefore, we will refer to mCherry/*CLV3*-positive nuclei as stem cells and mCherry/*CLV3*-negative samples from the closest neighboring tissue as non-stem cells.

**Figure 1.**
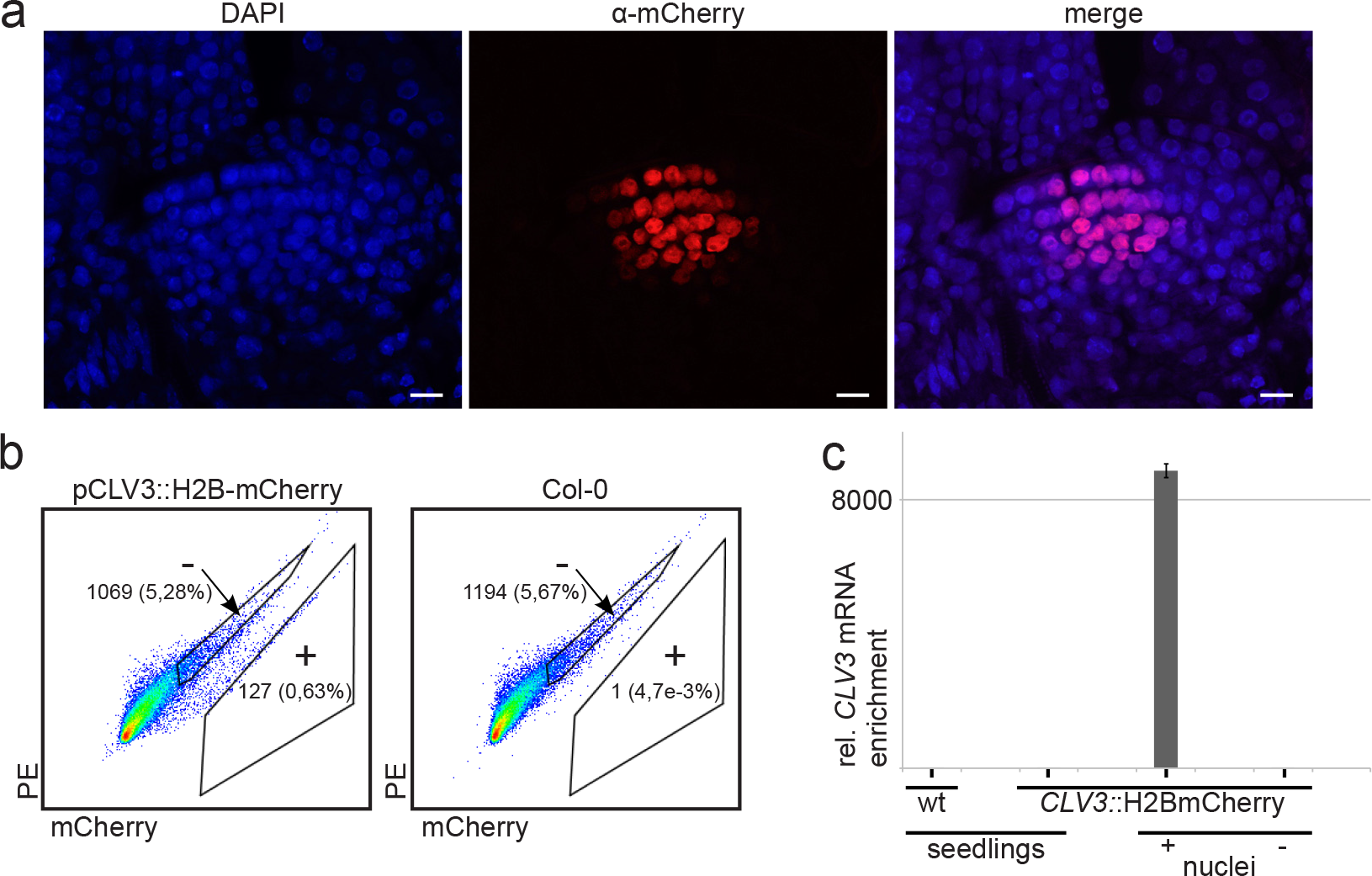
Establishment of FANS for stem cells of the shoot apical meristem (SAM). (**a**) Expression of H2B-mCherry under control of the *CLV3* promoter in 14 d-old seedlings. Whole-mount immunostaining using α-mCherry antibodies and laser scanning microscopy (scale bar 10 μm). (**b**) Example of a FANS experiment: mCherry-positive (+) and mCherry-negative (-) gates of DAPI-gated nuclei. Numbers indicate total number and percent of DAPI events. (**c**) Enrichment of *CLV3* transcript in mCherry-positive nuclei determined by qRT-PCR.

We generated and sequenced RNA expression libraries from stem and non-stem cells isolated from heart through torpedo stage embryos (E), 7 day- (D7), 14 day- (D14), and 35 day- (D35) old plants (Fig. S2a, b, Table S2), to (i) identify expression signatures in stem cells preceding major developmental switches, (ii) find genes that are involved in epigenetic resetting and germline formation, and (iii) detect “stemness” genes whose expression would characterize stem cells independent of development. Normalized read counts demonstrated high enrichment of *CLV3* and *mCherry* transcripts in stem cells at all developmental stages (Fig. 2a). High expression of the meristem marker genes *STM* and *KNAT1* relative to nuclei of 14-day-old whole seedlings (S14) confirmed the meristematic features of the non-stem cells (Fig. 2a).

**Figure 2.**
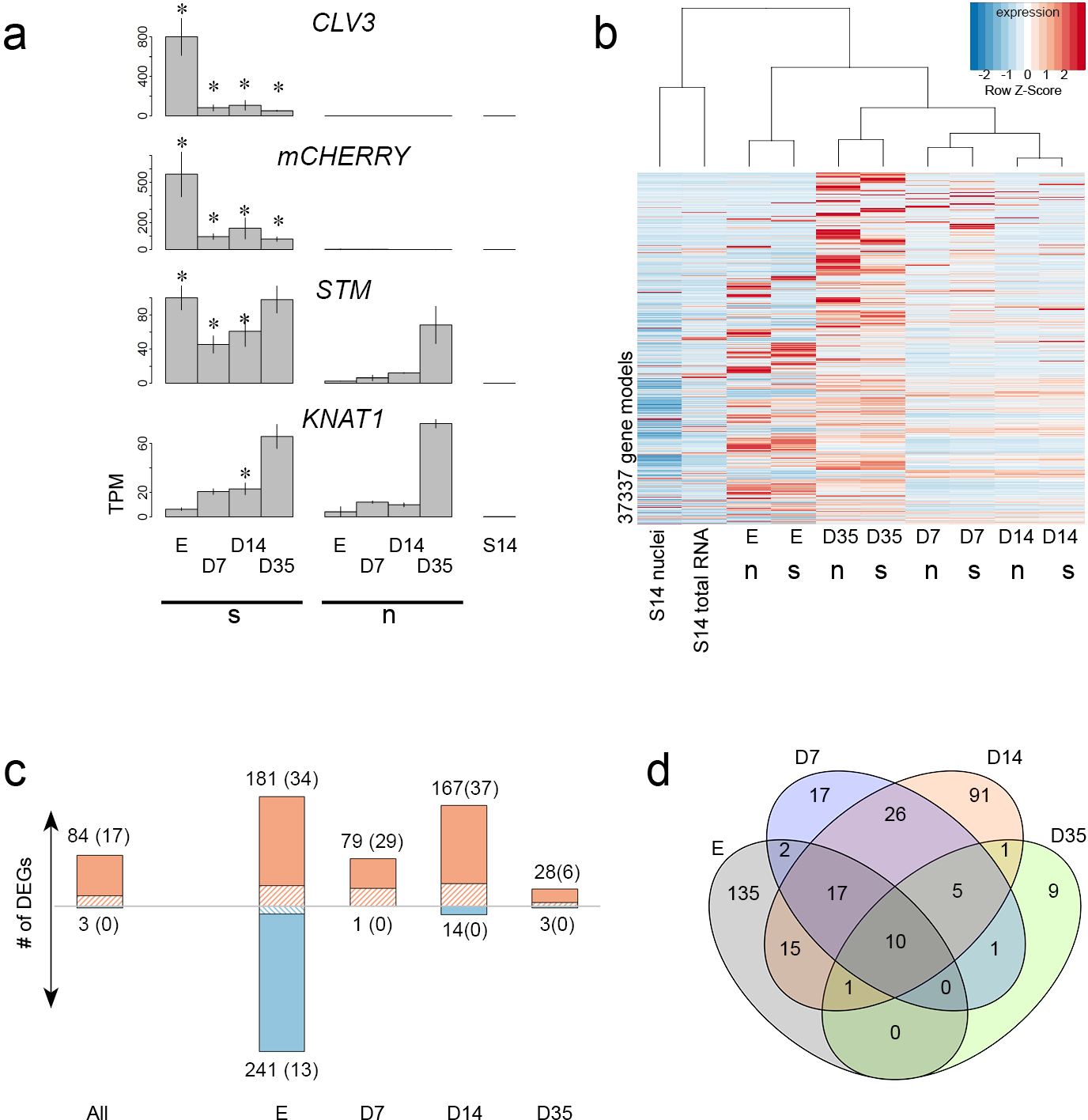
Differential RNA expression in SAM stem cells during development. (**a**) Expression of *CLV3*, *mCherry*, and the meristem marker genes *STM* and *KNAT1*. (**b**) Hierarchical clustering of expression data. (**c**) Number of DEGs between stem and non-stem cells at each timepoint. The banded portion of the bars indicates the number of transcription factor genes (also in parenthesis). (**d**) Overlap of genes with higher expression in stem cells (excluding *mCherry)*. s = stem cells; n = non-stem cells.

Transcriptome-wide clustering analysis showed that the expression signature of stem cells is dominated by developmental stage rather than cell type (Fig. 2b). Pairwise comparison between stem cells with the respective non-stem cells revealed differentially expressed genes (DEGs, q <0.05) at all timepoints (Fig. 2c), the majority upregulated in stem cells (with the exception of the embryo samples). GO term analysis revealed that GOs describing reproductive processes, floral organ development, and inflorescence development were already enriched in E, D7, and D14 (Table S3, Fig. S3), while their absence in D35 stem cells was likely due to the low number of DEGs.

Overlap analysis between samples (Fig. 2d, Fig. S4) revealed many stage-specific DEGs but also identified a set of 10 core genes (including *CLV3*) that were more highly expressed in stem cells of all four stages (Fig. 2a, Fig. S4 and S5, Table S4), and 23 genes with elevated expression in three out of the four stages (Table S4). Twelve of these 33 genes encode transcription factors (p-value for enrichment: 1.24e-08). Seven were previously connected with a meristem- or stem cell-related function (Table S4) leaving the remaining 26 as candidates with a potential role in stem cell maintenance.

We could not detect significant overlap with transcript analysis in the SAM during flower induction ^8^, probably due to differences in experimental set up and tissue type. The meristem transcriptome of the *ap1-1;cal1-1* double-mutant ^6^ had limited but significant overlap for upregulated genes (Table S5). Comparison with transcriptome data for different types of root meristem cells ^11^ resulted in an overlap especially with upregulated genes from *WOX5*- expressing cells of the quiescent center (Table S5). Also noticeable was an overlap between upregulated stem cell DEGs with genes related to DNA methylation or siRNAs highly expressed in meristematic tissue ^12^ (Fig. 3a). Among these are transcripts of two Argonaute proteins (AGO5 and AGO9), two histone methyltransferases (SUVH4 and SUVR2), the nucleosome remodeler DDM1, and three putative RNA-dependent RNA polymerases (RDR3, 4, and 5). This indicated that specific family members of prominent epigenetic components were upregulated in stem cells. Since AGO9, SUVH4, and DDM1 (among others) are necessary for TE repression ^13–15^, we asked whether TEs were downregulated in stem cells relative to the surrounding cells. Indeed, several Arabidopsis TE families ^13–16^ were 2-fold less expressed in stem versus non-stem cells through the four stages (Fig. 4 and Table S6). Surprisingly, with the same significance threshold, we found other TE families that were more highly expressed in stem cells (Fig. 4, Table S6). Strikingly, D7 showed the largest number of highly expressed groups and the lowest number of downregulated groups, indicating a transient loss of control over TE expression in stem cells at this early stage of vegetative growth, followed by resilencing towards generational transition.

**Figure 3.**
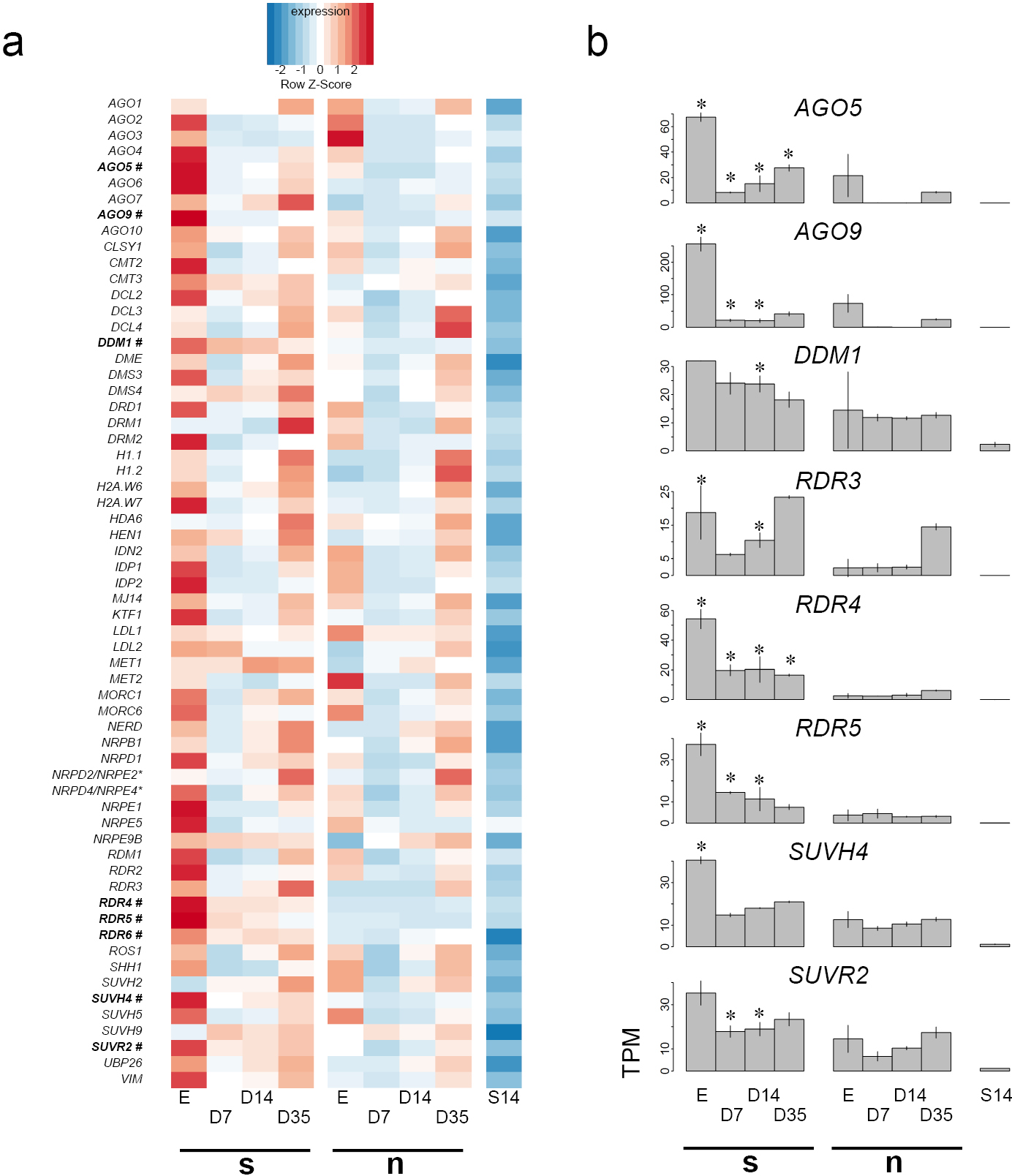
Expression analysis of genes related to epigenetic regulation. (**a**) Expression heatmap (in alphabetical order of gene acronyms). (**b**) Expression of significantly upregulated DNA methylation-related genes in stem cells, marked with # in (a). Asterisks indicate timepoints of significantly different expression between stem and non-stem cells. s = stem cells; n = non-stem cells.

**Figure 4.**
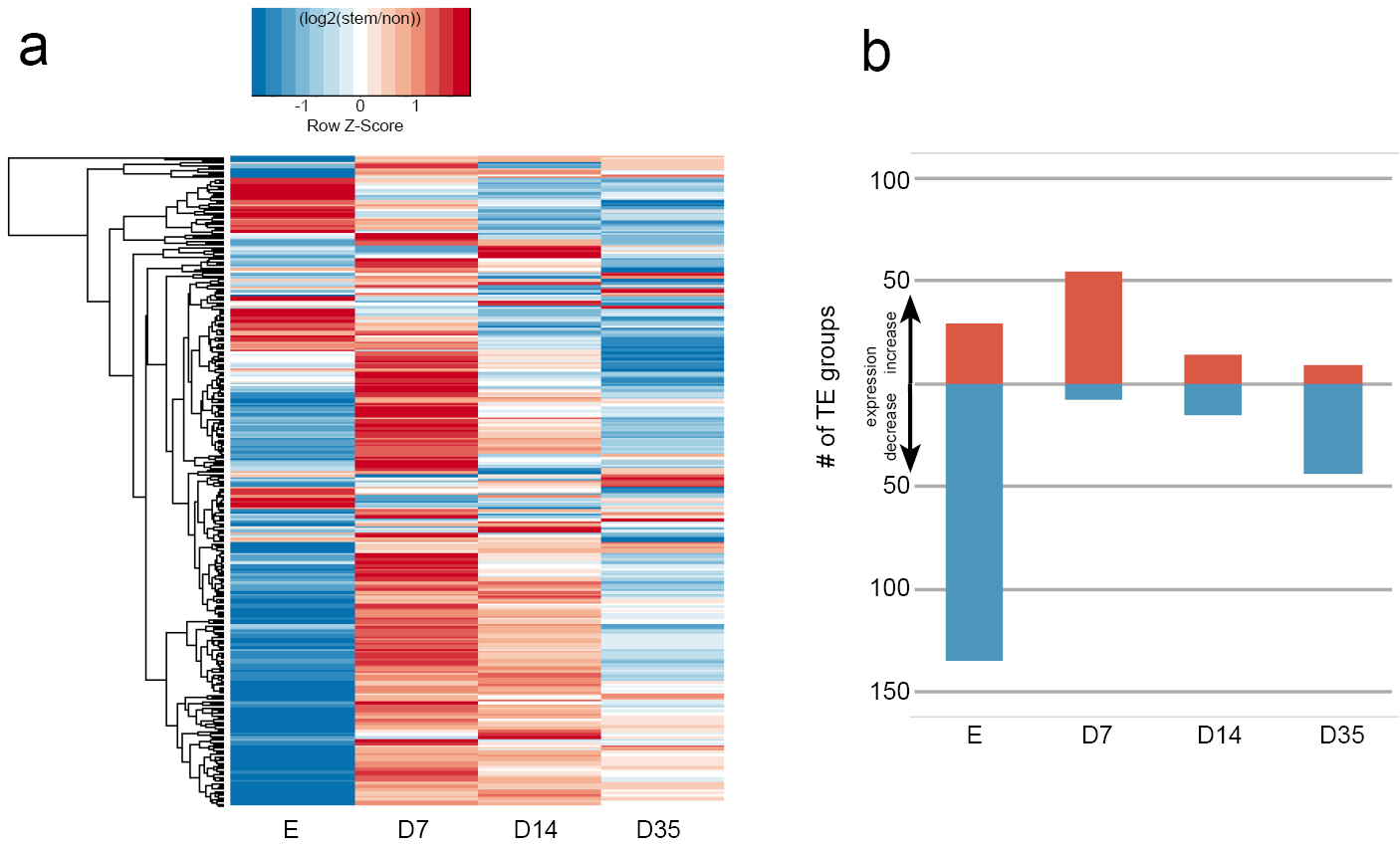
Expression analysis of transposable elements. (**a**) Heatmap of expression differences for all 318 Arabidopsis TE groups in stem cells relative to non-stem cells at different timepoints. (**b**) Number of TE groups with at least 2x expression difference at the different timepoints.

TEs overexpressed in D7 were mostly COPIA LTR-retroelements and Mutator-like DNA transposons but also included Helitrons, gypsy-like LTR elements, and SINEs (Table S6). As LTR retroelements are more prevalent within pericentromeric regions, whereas SINEs and Helitrons are distributed on chromosome arms ^17^, TE expression in stem cells occurred independently of chromosomal localization. We could not find a bias for TEs that were recently mobile in natural populations ^18^, nor for transposons with new insertions in DNA methylation-deficient mutants ^19, 20^.

To determine whether TE expression was influenced by changing DNA methylation, we performed whole-genome bisulfite sequencing of genomic DNA from D7, D14, and D35 stem and non-stem nuclei, with material from 7 d- and 14 d-old seedlings as reference. Modification of cytosines in plants (reviewed in ^21^) at CG sites (mCG) is mainly achieved by MET1 and occurs in repetitive sequences as well as along gene bodies. Cytosine methylation at CHG sites (mCHG) (H = A, C, or T) is installed by CMT2 and CMT3 and at CHH sites (mCHH) by DRM1 and DRM2 as well as CMT2. mCHG and mCHH are mostly restricted to repetitive sequences and important for TE silencing. mCHG is recognized by the histone methyltransferase SUVH4 which methylates histone H3 on lysine 9, a binding site for CMT3, and thereby reinforces DNA methylation in heterochromatic domains ^22^.

Analysis of DNA methylation distribution revealed pronounced differences around the centromeres for mCHG and mCHH, with the highest mCHG and lowest mCHH portion in stem cells of D35 (Fig. 5a and Fig. S6). Congruent with the distribution along the chromosomes, metaplot analyses revealed that these methylation differences were found at TEs, while protein-coding genes were not affected (Fig. 5b). mCHG levels increased with developmental age, and TEs in stem cells had consistently higher mCHG levels than the respective non-stem cells, reaching a maximum at D35. Conversely, mCHH decreased with developmental age, most pronounced in stem cells (Fig. 5b).

**Figure 5.**
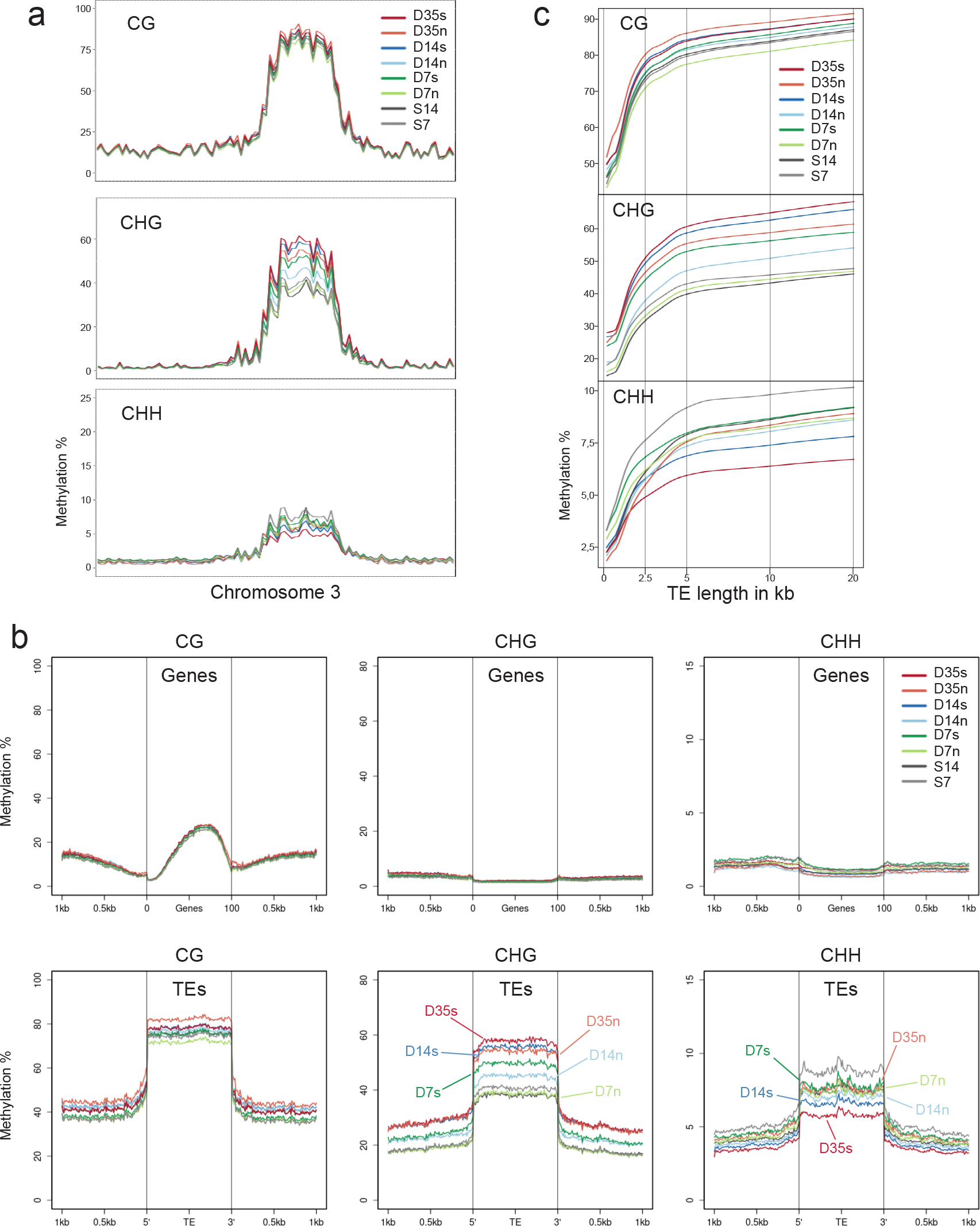
DNA methylation analysis of stem cells at different developmental stages. (**a**) CG, CHG, and CHH methylation at chromosome 3 in stem and non-stem cells. (**b**) Metaplots of DNA methylation at CG, CHG, and CHH for genes and transposons. (**c**) Locally weighted scatterplot smoothing fit of CG, CHG, and CHH methylation levels in stem cells and non-stem cells plotted on TE size. D7, D14 and D35 = sorted nuclei 7, 14, and 35 d.a.g., S7 and S14 = above-ground seedlings 7 and 14 d. a. g., s = stem cells; n = non-stem cells.

While TE groups varying in genomic location, cytosine content, structure and localization of repeats, and siRNA targeting sites ^17^ showed similarly increasing mCHG and decreasing mCHH in stem cells over developmental time (Fig. S7), there was a correlation with their length: plotting methylation levels of TEs against their size range (Fig. 5c) revealed that mCHG in older meristems increased more in long TEs (>2.5 kb), parallel to decreasing mCHH. This suggests a contribution of DDM1, as it mediates methylation preferentially at long TEs ^23^.

To understand which DNA methylation components are involved in methylation dynamics in stem cells, we identified differentially methylated regions (DMRs) for each timepoint and compared them with DMRs of mutants lacking different epigenetic components ^24^. Increased mCHG in stem cells was especially pronounced at D14 in hypo-DMRs of *ddm1*, *suvh4*, *cmt3*, and *suvh456;* DMRs with reduced mCHH overlapped with those of *cmt2*, *suvh456*, *ddm1*, and *met1* (Fig. 6). This suggests a concerted action between DDM1 and the reinforcing heterochromatin formation of CMT3 and histone methyltransferases to establish strong CHG methylation in stem cells entering the reproductive phase. Furthermore, the reduction of CHH methylation in *cmt2* and *suvh456* DMRs indicated a functional interference of CMT3 activity in stem cells with the related CMT2.

**Figure 6.**
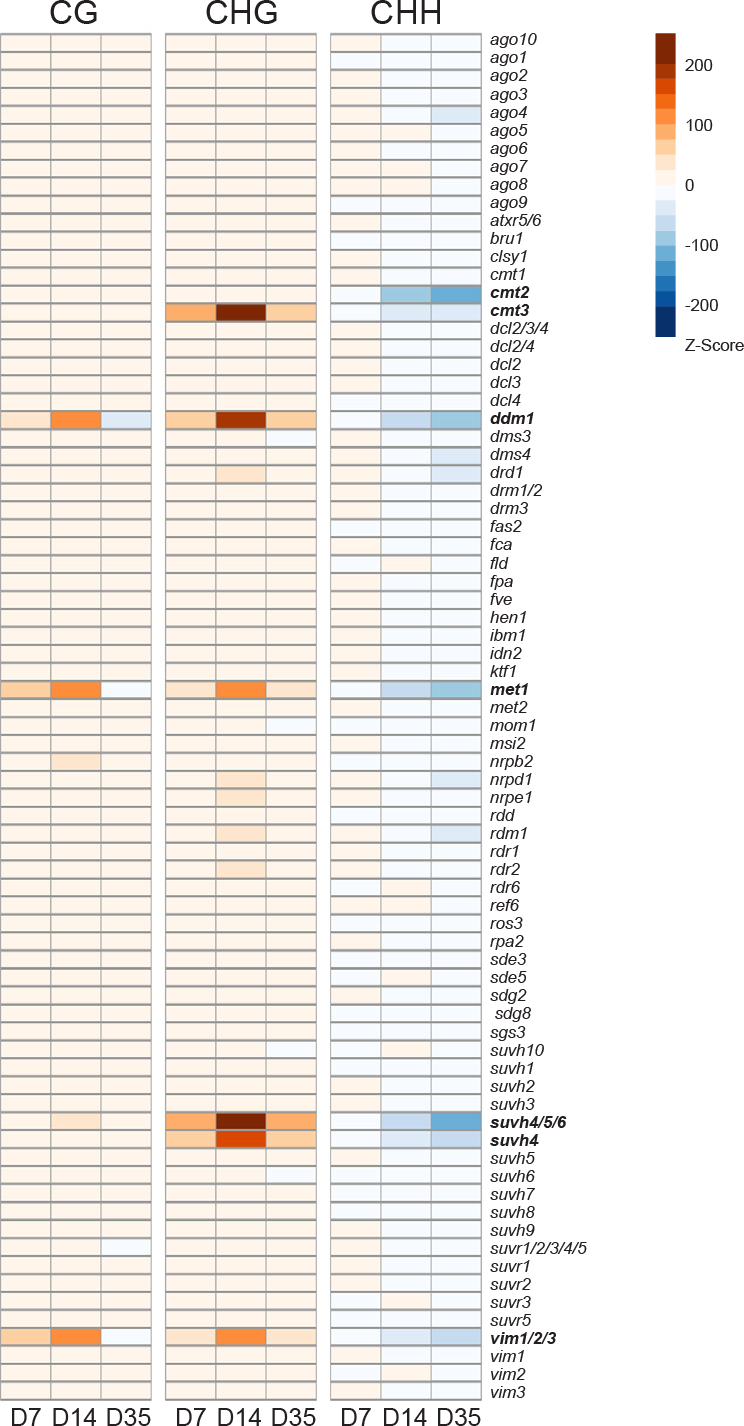
DMR analysis of stem cells. DNA methylation differences between stem and non-stem cell nuclei within DMRs of different epigenetic mutants. For each category, the scale denotes the number of standard deviations of differential methylation in relation to the rest of the genome.

The elevated TE expression at D7 correlated with a minimum of *AGO5* and *AGO9* expression (Fig. 3b). While they belong to different clusters of the AGO clade ^25^, both were shown to be expressed in meristematic tissue of embryos ^26, 27^ or in gametes or gametophytes ^28, 29^. AGO5 has not been connected with RNA-directed DNA methylation (RdDM) of TEs ^30^, and neither *ago5* nor *ago9* showed many DMRs in DNA of whole seedlings ^24^. However, AGO9 can restore methylation in an *ago4* mutant if accordingly expressed ^26^, suggesting that it can substitute RdDM-related functions. Although the specific molecular functions of these AGOs and the subpopulation of bound small RNAs in stem cells remain to be determined, their expression anticorrelated with active TEs could hint to a specific protection of germline precursor cells from virus and/or TE invasions. The transient loss of TE control in early vegetative stages might even provide the sequence-specific information, via small RNAs, for a stem cell-enriched or - specific silencing machinery at later stages. Mutants lacking major components of the RNA-directed DNA methylation (RdDM) pathway have no, or only mild, developmental defects. Reinforced silencing in stem cells during development, involving additional specific factors like RDR3, 4, or 5, may be responsible for this resilience. Such resilience might break down under special conditions, as indicated by stress-induced transposition prior to flower formation that only occurs in RdDM-compromised mutants ^19^. Interestingly, a recent study showed that male premeiotic meiocytes also exhibit high mCG and mCHG methylation and low mCHH ^31^. This raises the intriguing possibility that SAM stem cells enter a germline DNA methylation state long before they can be cytologically distinguished. Alternatively, our data could also suggest the presence of several cell types within the central domain of the SAM. The possibility to extend the isolation of stem cells at different stages, from mutants and under different environmental conditions, will enable future experiments to shed more light on epigenetic maintenance and dynamics in germline precursor cells.

## Data access

DNA bisulfite and RNA-seq data have been deposited in the ArrayExpress database at EMBL-EBI (www.ebi.ac.uk/arrayexpress) under accession number E-MTAB-5478 and E-MTAB-5479.

## Methods

### Plant material

All experiments were performed with *Arabidopsis thaliana* ecotype Col-0, wild type or transgenic for p*CLV3:H2B-mCherry*. The pCLV3:H2B-mCherry construct was generated as follows: the coding sequence of the H2B gene was PCR-amplified with primer H2B-forward and H2B-reverse (Table S7) from cDNA prepared from 14 d-old seedlings. The vector pCLV3:erCFP ^9^ was cut with *BamH*I and *Sac*I, and the H2B amplicon was inserted (In-Fusion, Clontech) into the open vector. The resulting plasmid was opened with *Sac*I and In-Fusion-filled with a PCR-amplified mCherry-coding fragment using the primers mCherry-fusion-F1 and mCherry-fusion-R1 (Table S7). Correct sequence of the resulting vector pCLV3:H2B-mCherry was confirmed by Sanger-sequencing. The construct was used to generate transgenic plants by the floral dip method ^32^. Primary transformants were selected with glufosinate (Merck) and their progeny screened for lines with a segregation ratio of 3 resistant to 1 sensitive plant. Homozygous offspring were propagated for seed amplification.

### Growth conditions

All plants were grown either *in vitro* on GM medium with or without selection or on soil under a 16 h light/8 h dark regime at 21°C. Material was always harvested at the same time of the light period.

### Microscopic analysis and immunostaining

For wide-field microscopy, plant material was immersed in PBS buffer and imaged with a Zeiss Axio Imager epifluorescence microscope. Isolated nuclei were imaged with an LSM780 Axio Observer, and Images were deconvolved using Hyugens Core (Scientific Volume Imaging) with a theoretical PSF. Immunostaining was performed according to ^33^, with an additional clearing step using ScaleA ^34^ and DAPI as counterstain. Anti-mCherry nanobodies were purchased from Chromotek (#rba594-100). Immunostains of meristems were imaged using the Airyscan mode on an LSM880 Axio Observer.

### Fluorescence-activated nuclei sorting (FANS)

For 7D/14C/35D samples, 200-800 apexes (depending on size) of soil-grown plants with the corresponding age were collected. For embryo samples, ovules from siliques of a few representative plants were analyzed to contain early heart till early torpedo stage embryos, and developmentally identical siliques were used to dissect 3000-4000 ovules. Collected material was immediately transferred into nuclei isolation buffer on ice (NIB: 500 mM sucrose, 100 mM KCl, 10 mM Tris-HCl pH 9.5, 10 mM EDTA, 4 mM spermidine, 1 mM spermine and 0.1% v/v 2-mercaptoethanol, prepared just before use ^35^). The material was then transferred into a tube containing 1.8 ml of nuclear extraction buffer (NEB of the Sysmex CyStain^®^ PI Absolute P kit (#05-5022) plus 1% v/v 2-mercaptoethanol) and disrupted with the TissueRuptor (Qiagen) at the lowest speed for 1 min. The suspension was filtered (30 μm filter nylon mesh, Sysmex # 04-0042-2316) and centrifuged for 10 min at 4000 rcf at 4°C. The nuclear pellet was resuspended in Precise P staining buffer (Sysmex #05-5022; plus 1% v/v 2-mercaptoethanol and DAPI to a final concentration of 5 μg/ul), incubated for 15 min and again filtered (30 μm) into tubes (Sarstedt #55.484.001). Sorting was performed on a BD FACSAriaTM III cell sorter (70 μm nozzle). Forward/Side scatter and DAPI and mCherry gating were adjusted with wild type nuclei (DAPI-positive, mCherry-negative) as reference. The mCherry gate was adjusted so that a maximum of 1/10 of mCherry events occurred in wild type compared to the p*CLV3:mCherry-H2B* line. For DNA extraction, nuclei were directly sorted into Genomic Lysis Buffer (Quick-DNA Microprep Kit, Zymo Research, #D3020,), and DNA was purified according to the suppliers’ protocol for whole blood and serum samples. DNA was quantified using pico-green on a NanoDrop fluorospectrometer (Thermo Scientific). For RNA isolation, NIB, NEB, and staining buffer were complemented with RiboLock RNase inhibitor (Thermo Scientific #EO0381, final concentration 1 U/μl) and nuclei were directly sorted into TRIzol LS (Ambion, #10296028). RNA was prepared according to the manufacturers’ recommendation, except that nuclease-free glycogen (Thermo Scientific) was added during an overnight precipitation at −20°C. Amount and quality of RNA was determined on an RNA 6000 pico-chip (Bioanalyzer/Agilent Technologies). For DNA and RNA extraction, DNA-LoBind tubes (Eppendorf, #022431021) were used.

### qPCR analysis

For qPCR and enrichment analysis, RNA was extracted with TRIzol LS (Ambion) either from sorted nuclei or from shock-frozen and ground tissue material. RNA was treated with DNAse (Thermo Scientific, #79254) and reverse-transcribed with iScript (Biorad, #172-5038). qPCR assays were performed with Universal ProbeLibrary (UPL) assays (Roche, # 06402682001) with primers and probes described in Table S7.

### Library preparation and sequencing

For RNA library preparation, total RNA of biological duplicates was extracted either from nuclei directly sorted into TRIzol LS or from shock-frozen ground material and used to generate cDNA libraries with the SMART-Seq v4 Ultra Low Input RNA Kit (Clontech). For the comparison with the nuclear RNA transcriptome, RNA was extracted from DAPI-stained FANSed nuclei isolated from 14 d-old p*CLV3:mCherry-H2B* seedlings with the same protocol as for cDNA production. cDNA populations were paired-end sequenced on a HiSeq 2500 Illumina sequencing platform. For bisulfite library preparation, at least 200 pg of DNA was used. Libraries were prepared with the Pico Methyl-Seq Library Prep Kit (Zymo Research #D5456) according to the manufacturer’s protocol.

### Analysis of the RNA-sequencing data

For the analysis of nuclear to total RNA expression correlation, Tophat ^36^ was used for mapping to the TAIR10 reference genome after removal of low-quality bases with Trimmomatic ^37^ (parameters: LEADING:8 TRAILING:10 SLIDINGWINDOW:4:15 MINLEN:50;). Cuffdiff ^38^ was used for normalization.

For all other analysis, RNA-seq reads were first adapter- and quality-trimmed with Trim Galore! (Krueger F. Trim galore, v0.4.1, with default parameters). The reads were then aligned to the TAIR10 reference genome (including the mCherry sequence) with STAR ^39^ (v2.5.2a) (Table S8). Alignment parameters for STAR were set by the quantifier RSEM ^40^ (v1.2.30), which are based on previous ENCODE standards. The annotation used for quantification was Araport11. RSEM was run with default settings. To correct for possible positional biases in the data, we activated RSEM’s positional bias correction option (–estimate-rspd). The resulting gene expression tables were imported into R (v3.4) via the tximport package ^41^ (v1.4.0). Consecutive differential gene expression analysis was performed with DESeq2 ^42^ (v1.16). Samples of the same stages were analyzed pairwise via DESeq2s Wald test (FDR < 0.05). To detect genes that are differentially regulated in stem cells across all timepoints, we made use of DESeq2’s model-based likelihood ratio test (LRT, FDR < 0.05). The LRT allowed us to investigate how well the expression of a gene is recapitulated by different models. DESeq2 compares two models, one full model and a reduced model. Our full model factored in the cell type, the stage, and the interaction of both, while our reduced model did not factor in the interaction, leaving us with a set of differentially expressed genes whose variation can be explained by a combination of cell type and time. The RNA-seq pipeline is available under https://gitlab.com/nodine-lab/rsem-rna-seq-pipeline/. GO enrichments were calculated using the AmiGO2 tool and the PANTHER classification system (http://amigo.geneontology.org/rte) ^43^. Visualization and clustering of the data was achieved using the R packages “gplots” and “gclus”.

### DEG TE-Families

All RNAseq samples were quality-trimmed using cutadap (v1.14) (Marcel Martin; Cutadapt removes adapter sequences from high-throughput sequencing reads; EMBnet.journal; Vol17, No1) and trimmomatic ^37^ (v0.36). STAR ^39^ (v2.5.2a) (Col-0 Arabidopsis reference genome, the Araport11 gene and TE annotations) was used as reference to map the reads, allowing multiple hits (–outFilterMultimapNmax 100 and --winAnchorMultimapNmax 100). TEtranscripts from the TEToolkit ^44^ (v1.5.1) was used in multi-mode to find DEG TE-families.

### Analysis of the bisulfite-sequencing data

Illumina HiSeq 2500 sequencing data was obtained from three stages (D7, D14, and D35) each in three different settings (+: FANS-sorted stem cell tissue, -: non-stem cell but meristematic tissue, s; whole seedling). Samples D14 and D35 were sequenced with 125 bp paired end reads, D7 with 50 bp paired end reads (Table S8). The data were quality-checked (fastqc) and trimmed with TrimGalore (Krueger F. Trim galore, v0.4.1, default settings with stringency = 1) and trimmomatic ^37^ (v0.36, sliding window: 4:20, leading: 20). Bismark ^45^ (v0.18.1 with Bowtie2 v2.2.9) was used to map the reads to the *Arabidopsis thaliana* Col-0 reference genome (including mitochondria and chloroplast genomes) in the non-directional mode with a mapping stringency of L,0,-0.6. A mapping-position-based removal of duplicates (Bismark) was applied, and the C-to-T conversion rate was calculated using the reads mapped to the chloroplast genome (ranging from 98.9 to 99.5%). Methylation was called (Bismark), ignoring the first bases according to the M-Bias plots. Samples with same stages and settings were pooled to a single sample, resulting in genome coverages for the nuclear genome from 16,4x to 53,9x.

### DMR analysis

Differentially methylation positions (DMP) were identified by Fisher’s exact test. Their positions were clustered together based on a minimum distance of 50 bp between DMPs to call a differential methylated region (DMR). DMR calling was done using methylpy (https://github.com/yupenghe/methylpy.git) version 1.1.9. We used custom R and python scripts for further analysis of these DMRs.

## Author contributions

R.G. and O.M.S. designed the study and wrote the manuscript, R.G., N.D., A.G., G.B., N.L., and M.D. performed the experiments, K.R., T.N., R.P., and F.H. analyzed the data, Ma.N. and Mi.N. discussed the results and commented on the manuscript.

## Acknowledgments

We are grateful for excellent support by the GMI/IMP/IMBA Biooptics facilities, the Next Generation Sequencing and Plant Sciences units of the Vienna BioCenter Core Facilities (VBCF). We thank Claude Becker, Frederic Berger, and James Matthew Watson (all GMI, Vienna) for helpful comments on the manuscript, and Thomas Laux for the CLV3 promoter. We gratefully acknowledge financial support from the Austrian Science Fund (FWF I489, I1477 to O.M.S and I3687 to R.G.) and the Plant Fellows program (EU FP7) to R.G.

## Competing interests

The authors declare no competing financial interest.

---

**Supplementary Figure 1:**
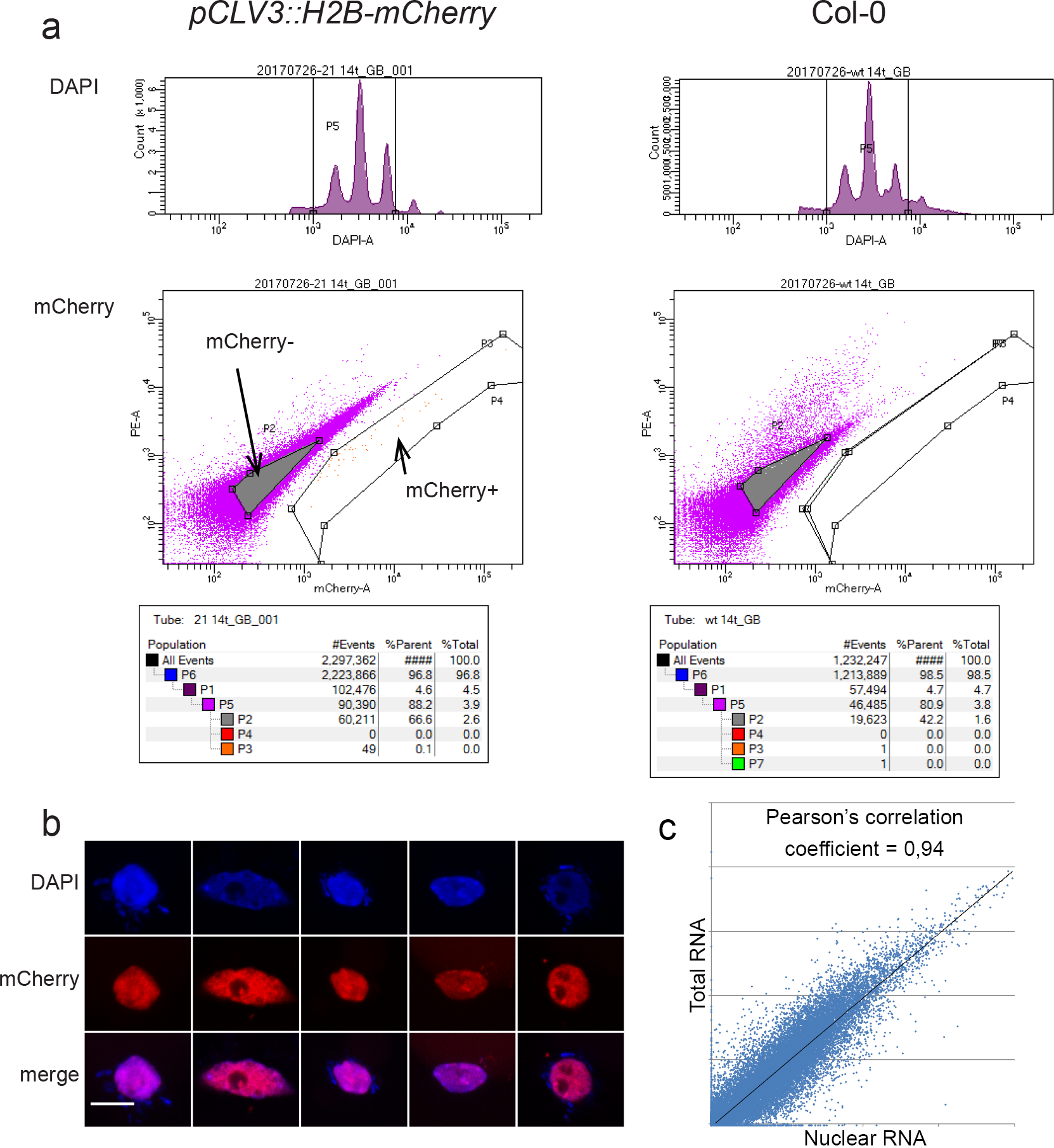
Isolation of stem cell nuclei and RNA comparison. (**a**) Gating strategy used for FANS of stem cells. Representative FANS plots are shown. Events are gated for DAPI (top row) and next either for mCherry+ or mCherry-(bottom row). For numbers see also Table S1. (**b**) Examples of mCherry-positive nuclei after FANS (scale bar 5 μm). (**c**) Correlation of log10-normalized FPKM values of nuclear and total RNA extracted from 14 d-old seedlings.

**Supplementary Figure 2:**
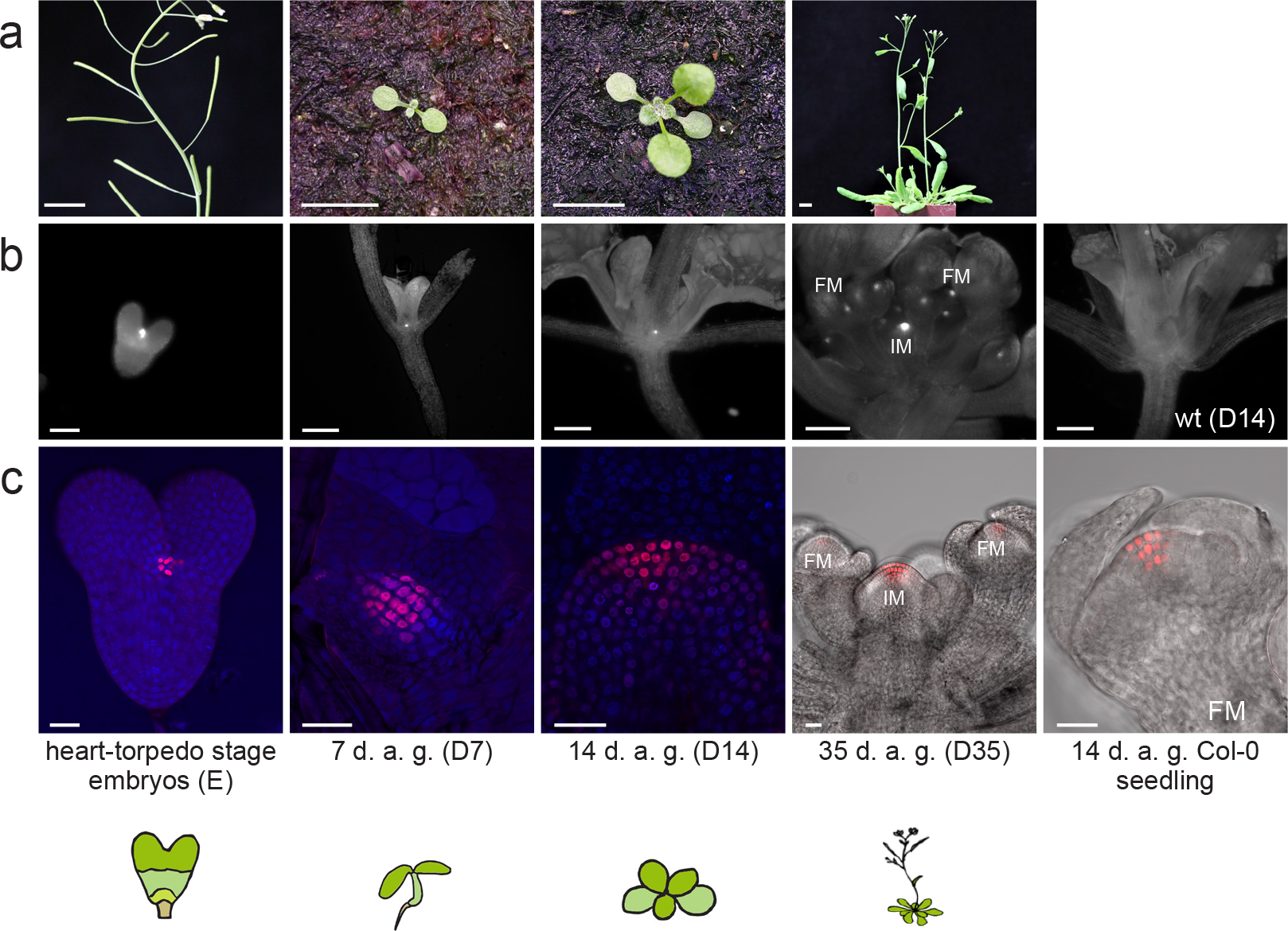
Growth stages used for genome-wide expression and DNA methylation analysis in stem and non-stem cells. (**a**) Developmental stages of representative plants (scale bars 1 cm). (**b**) Wide-field microscopic images with RFP filters. (**c**) LSM pictures of representative plants. For better visualization DAPI was used as counterstain in E, D7 and D14. IM = Inflorescence meristem. FM = Floral meristem. Scale bars in (b): 60 μm for the embryo; 1 mm for the other three stages. Scale bars in (c): 20 μm.

**Supplementary Figure 3:**
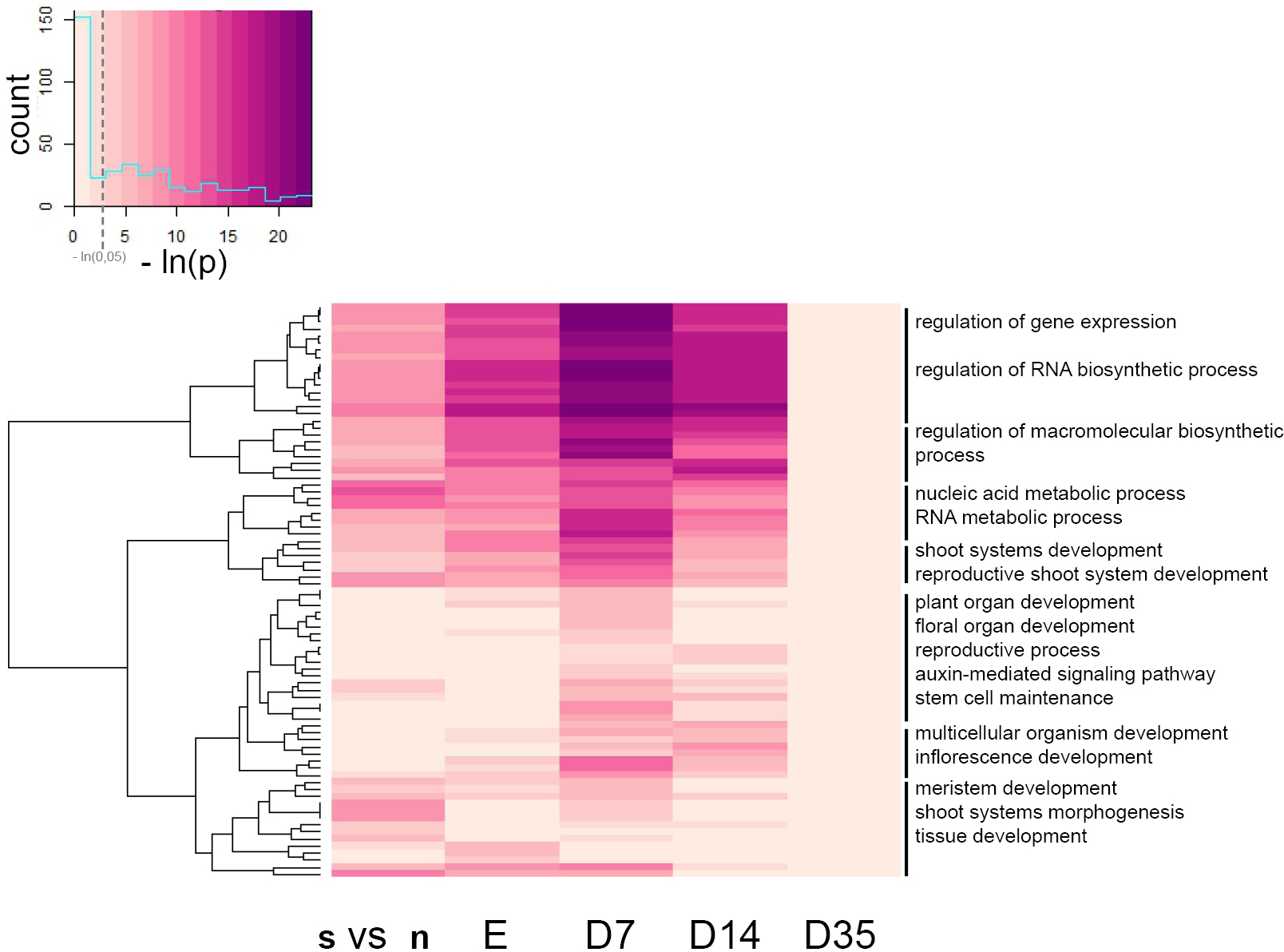
Clustered heatmap displaying GO-term enrichment. Color codes represent the negative ln of the Bonferroni corrected p-value for enrichment of each GO-term. A p-value of 0.05 corresponds approximately to 3. See also Table S3 for exact values. s = stem cells; n = non-stem cells.

**Supplementary Figure 4:**
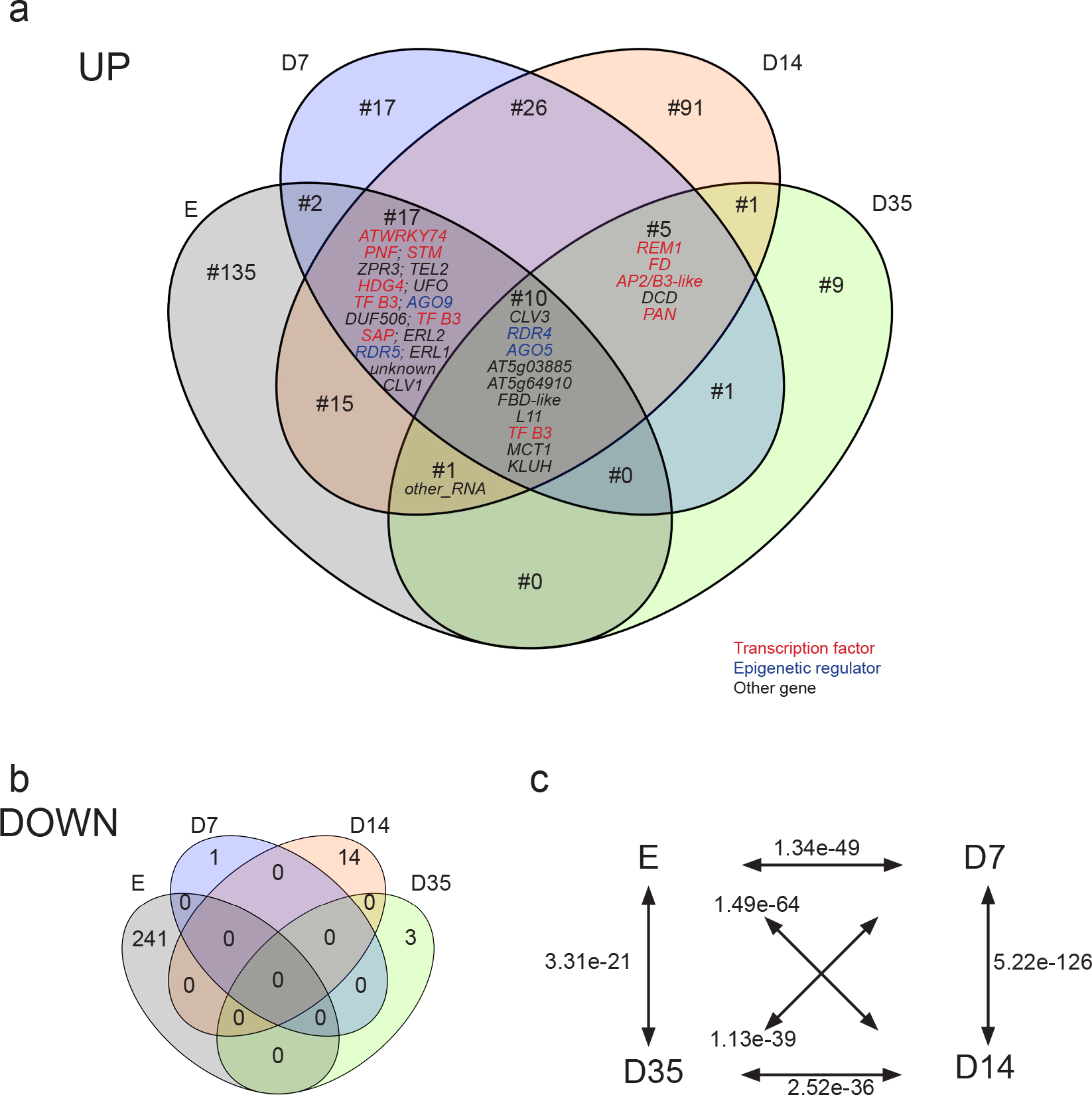
Overlap of DEGs at different timepoints. (**a**) Venn diagrams for genes up- and (**b**) downregulated in stem cells, respectively. (**c**) p-values (hypergeometric tests) for likelihood of overlap of upregulated genes in different pairs of timepoints.

**Supplementary Figure 5:**
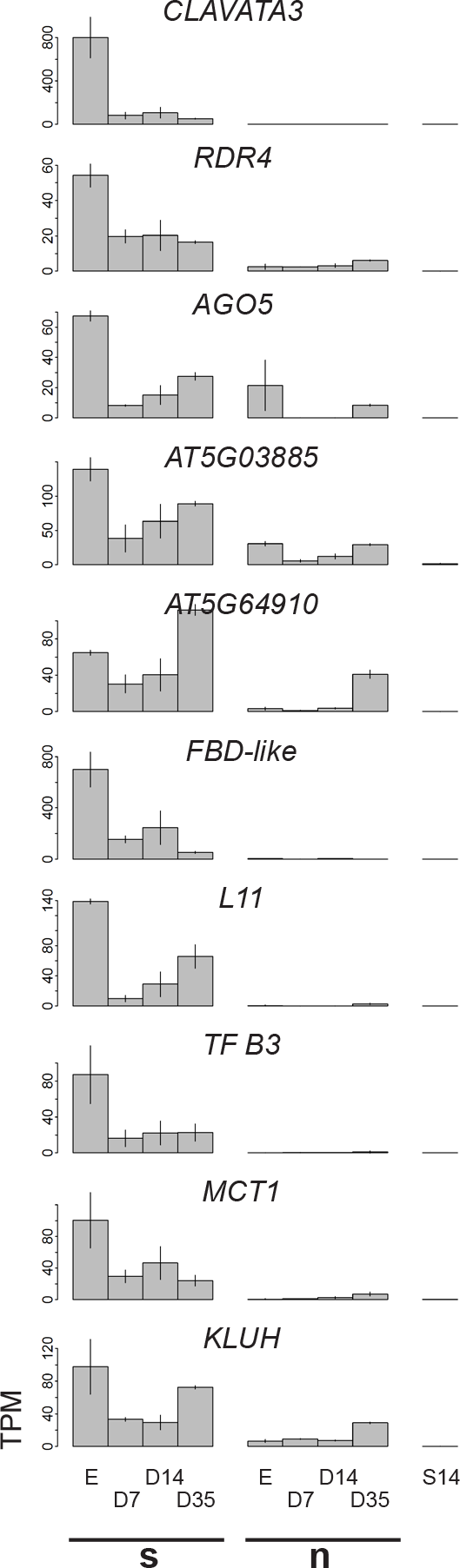
Expression of core stemness genes. Bar plots of expression of genes that are significantly upregulated in SAM stem cells throughout development. s = stem cells; n = non-stem cells.

**Supplementary Figure 6:**
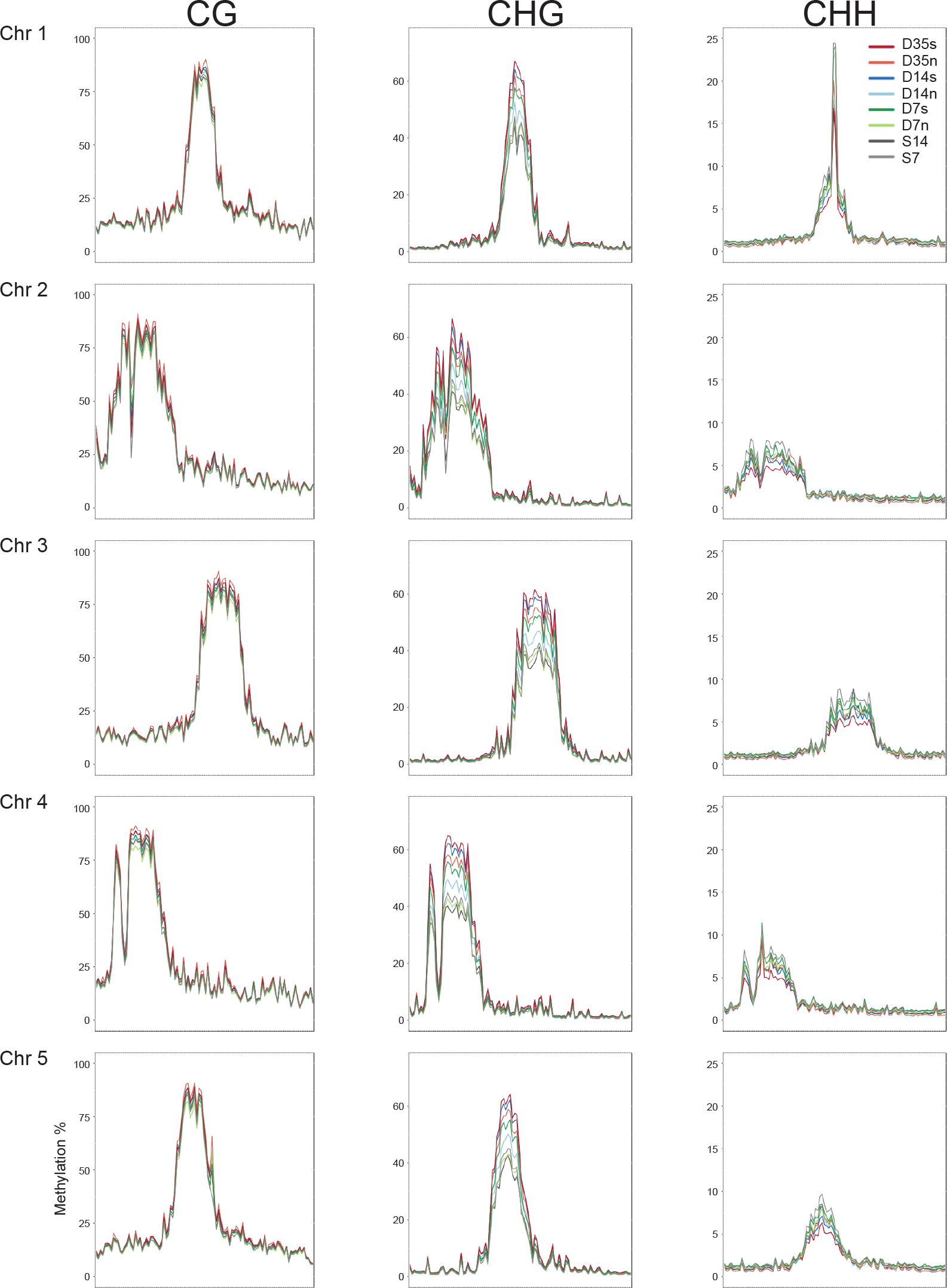
DNA methylation analysis of stem cells on all five Arabidopsis chromosomes in stem and non-stem cells at different developmental stages. s = stem cells; n = non-stem cells.

**Supplementary Figure 7:**
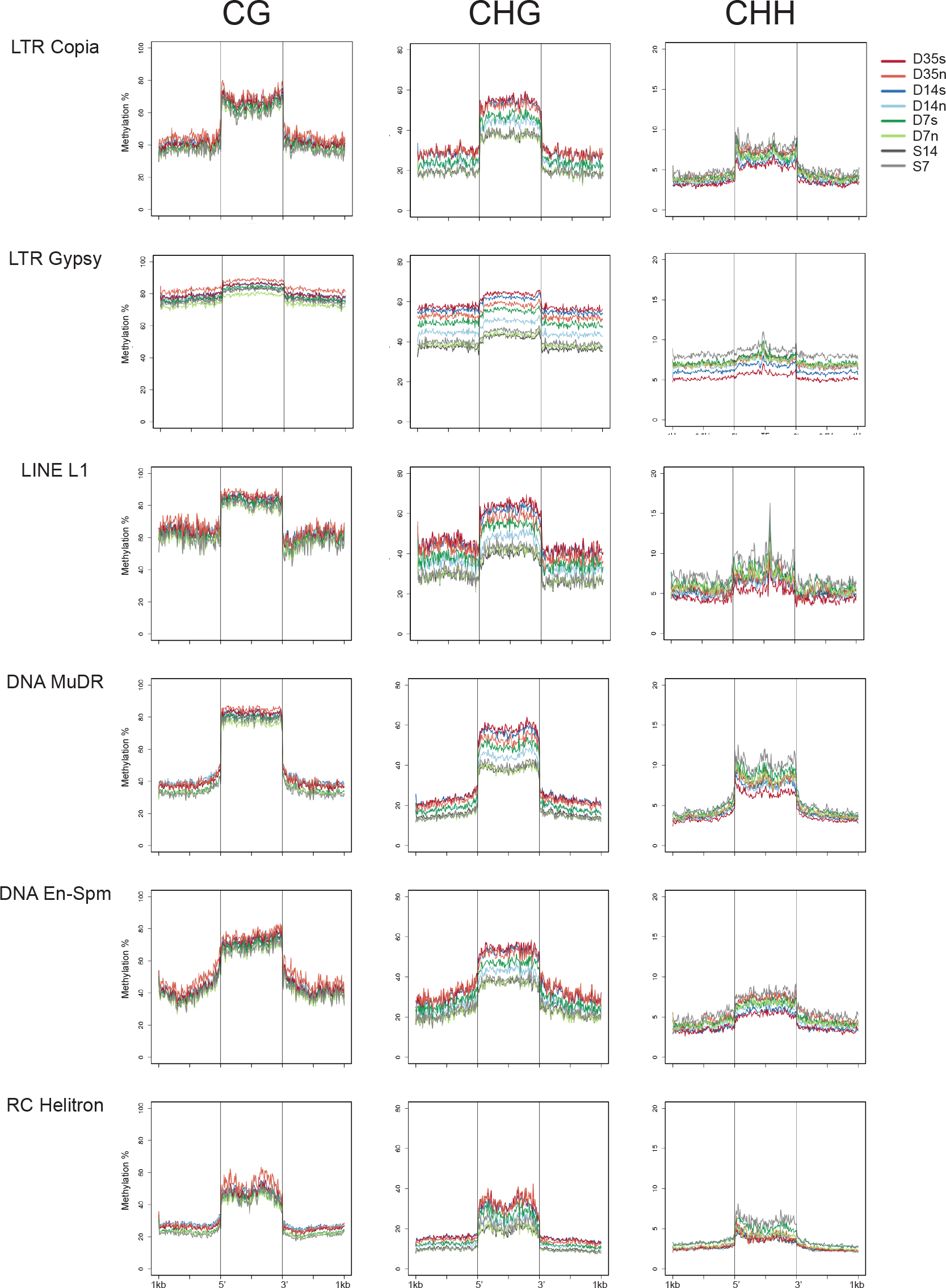
DNA methylation analysis of different TE classes in stem and non-stem cells at different developmental stages. s = stem cells; n = non-stem cells.

**Supplementary Table 1: Examples of FANS data**

**Supplementary Table 2: RNA expression data**

**Supplementary Table 3: GO-term annotations**

**Supplementary Table 4: Overlapping DEGs**

**Supplementary Table 5: Comparison with other data sets**

**Supplementary Table 6: TE expression data Supplementary Table 7: Primer sequences**

**Supplementary Table 8: Mapping statistics**

